# The effect of Dicer knockout on RNA interference using various Dicer substrate interfering RNA structures

**DOI:** 10.1101/2020.04.19.049817

**Authors:** Min-Sun Song, John J Rossi

## Abstract

Dicer-substrate siRNA (DsiRNA) was a useful tool for sequence-specific gene silencing. DsiRNA was proposed to have increased efficacy via RNAi gene silencing, but the molecular mechanism underlying the increased efficacy is not precise. We designed the tetra-looped DsiRNA as the tetra-looped RNAs have been reported more stable structure and increased binding efficiency with RNA and protein. To gain a deeper understanding of the Dicer function of DsiRNA, we knocked out Dicer in the HCT116 cell line and analyzed the efficacy of various Dicer substrates on RNAi gene silencing activity. Tetra-looped DsiRNA demonstrated increased efficacy of gene silencing Dicer expressing cells with activity favoring the guide strand. The gene silencing activity of all DsiRNAs was reduced in Dicer knockout cells. Thus, this study allows us to understand the Dicer function of key RNAi silencing and provides valuable resources for RNAi research and applications.

## Introduction

A hairpin structure is one of the most abundant RNA secondary structural elements. RNA hairpins play essential structural and functional roles by providing sites for RNA tertiary contacts and protein binding, which facilitates the assembly of ribonucleoprotein particles. More than half of all known RNA hairpins are composed of four nucleotides, called tetra-loops (1-5). Consequently, RNA tetra-loops are predominant and are commonly found in nature. Tetra-loops have been found in many RNAs, including ribosomal RNA (rRNA), mRNA, a group I intron ribozyme, RNase III, coxsackievirus B3, and heron Hepatitis B virus (6-10). In general, RNA tetra-loops are more stable compared to smaller or larger loops accommodating the same stem, because the compact structures of tetraloops impact them with high thermal stability and nuclease resistance.These loops involve base stacking, base-phosphate, and base-ribose hydrogen bonds (11). GNRA and UNCG (5’ to 3’, N: any base and R: purine) are the most common among the known stable tetraloops (12,13). Among these families, a GAAA tetra-loop has a U-turn motif, which can be involved in RNA activity by facilitating both RNA-RNA and RNA-protein interactions (14). It is noteworthy that RNA tetra-loops confer functional roles within the RNA beyond allowing secondary structure formation. The tetra-loops generally participate in RNA tertiary interactions with other RNAs and RNA-protein interactions (11,15).

Previously, chemically synthesized 25- to 27-nucleotide (nt)-long double-stranded RNAs (dsRNAs) with 2-nt 3’ overhangs were identified as substrates for the Dicer endoribonuclease. These Dicer-substrate siRNAs (DsiRNAs) are recognized and processed into shorter small interfering 21-22bp RNAs (siRNAs) by endogenous Dicer when they are introduced into mammalian cells (16). The interaction between DsiRNAs and Dicer promotes the loading of siRNAs into the RNA-induced silencing complex (RISC) with strand-specific orientation, and the association of the guide strand into the argonaute protein Ago2 in the cytoplasm (16-18) Subsequently DsiRNAs have been reported to be up to 10-fold more potent in silencing a targeted gene than canonical 21-nt siRNAs (19-21). In contrast, endogenous microRNA (miRNA) is processed from hairpin-containing primary transcripts (pri-miRNA) of nuclear hairpin double-strand RNA into pre-miRNA by the ribonuclease Drosha, before it is transported to the cytoplasm and further cleaved by Dicer (22). RNA containing hairpin structures influences Dicer’s activity and site selection (23). The loop structure of short hairpin RNAs (shRNAs), which are artificially designed small RNAs, contributes to the biogenesis and subsequent activity of siRNA with Dicer (24). How can the activity of small RNAs be improved act by Dicer? We may find the answer to this in the study of structures between Dicer and small RNAs. A cryo-electron microscopy structure study showed that the terminal loop of pre-miRNA interacts with the N-terminal DEAD-like helicases domain (DExD)/H-box helicase domain of Dicer and its co-factor, transactivation response element RNA-binding protein (TRBP) (25). This implies that the presence of loops within small RNAs may influence not only Dicer cleavage activity but also gene silencing efficacy.

In this study, we designed various DsiRNAs with or without a 5’-GAAA-3’ tetra-loop. To determine whether the multiple versions of DsiRNAs affected Dicer-dependent gene silencing efficacy, we generated Dicer knockout cells (named H2-2) in the colorectal cancer cell line HCT116. We compared the gene silencing activity of the various DsiRNAs in wild-type (WT) HCT116 and Dicer-inactivated H2-2 cells. Compared to WT cells, the gene silencing activity of DsiRNAs, including both original and tetra-looped versions, was reduced 2-fold to 37-fold in H2-2 cells. At present, it is not clear whether Dicer can directly affect the RNA interference (RNAi) activity of DsiRNAs. These results have important ramifications for the role of Dicer in the efficacy of DsiRNAs and tetra-looped DsiRNA biogenesis.

## Results

### Design of DsiRNAs targeting hnRNPH1

We previously observed successful gene silencing activity using DsiRNAs that targeted the RNA binding protein heterogeneous nuclear ribonucleoprotein H (hnRNPH1); this activity was associated with Ago2, TRBP, and Dicer (17,26). In the present study, we synthesized DsiRNAs specific to hnRNPH1 that also contained various tetra-loop and stem structures (Figure 1; Table 1). DsiRNAs that enhanced the efficiency of Dicer-mediated loading of siRNA into the RISC were designed as asymmetric duplexes containing a 27-base antisense strand with a 2-nt 3’-overhang and a 25-nt sense strand. Two DNA nucleotides were included at the 3’ end of the sense strand to create a blunt end (27). These asymmetric 25/27-mer siRNAs were optimized for processing by Dicer. We designed two versions of the DsiRNA (DsiRNA I [DI] and DsiRNA II [DII]), which differed by only a single base pair. Previously, our results showed that hnRNPH1-targeted DsiRNAs could show strand selectivity; DsiRNA I has similar selectivity for either strand, but DsiRNA II has more substantial activity on the antisense strand (17). We added a 5’-GAAA-3’ tetra-loop (TL) into each DsiRNA and synthesized different stem structures for each, one with a GC-rich stem (TL_DI and TL_DII), and another with the original stem (TL_DI_O and TL_DII_O), which corresponds to the hnRNPH1 target gene.

**Table 1.**
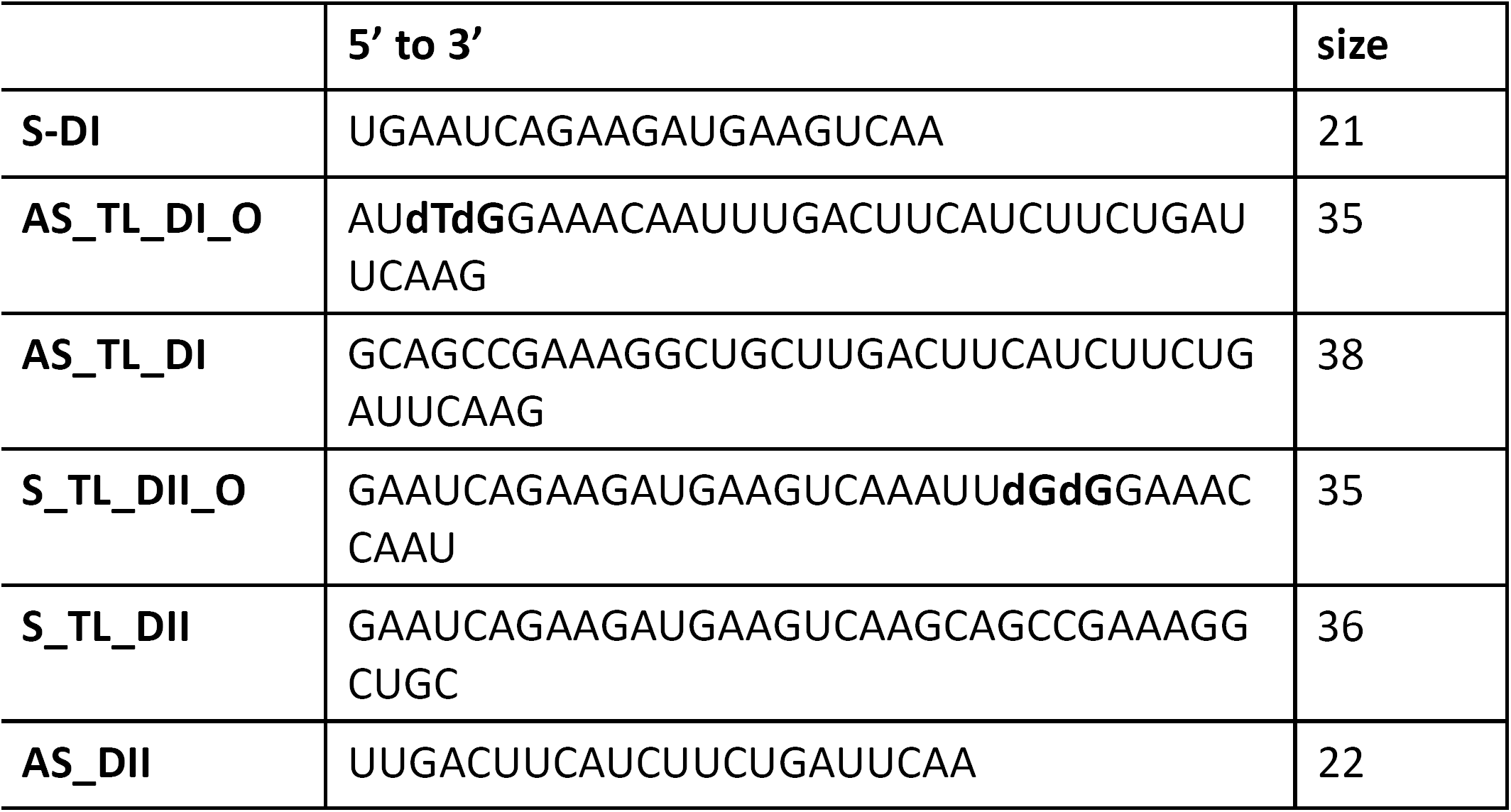
The sequence of each strand of DsiRNAs. DsiRNA sequences used to synthesize the tetra-looped DsiRNAs.

**Figure 1.**
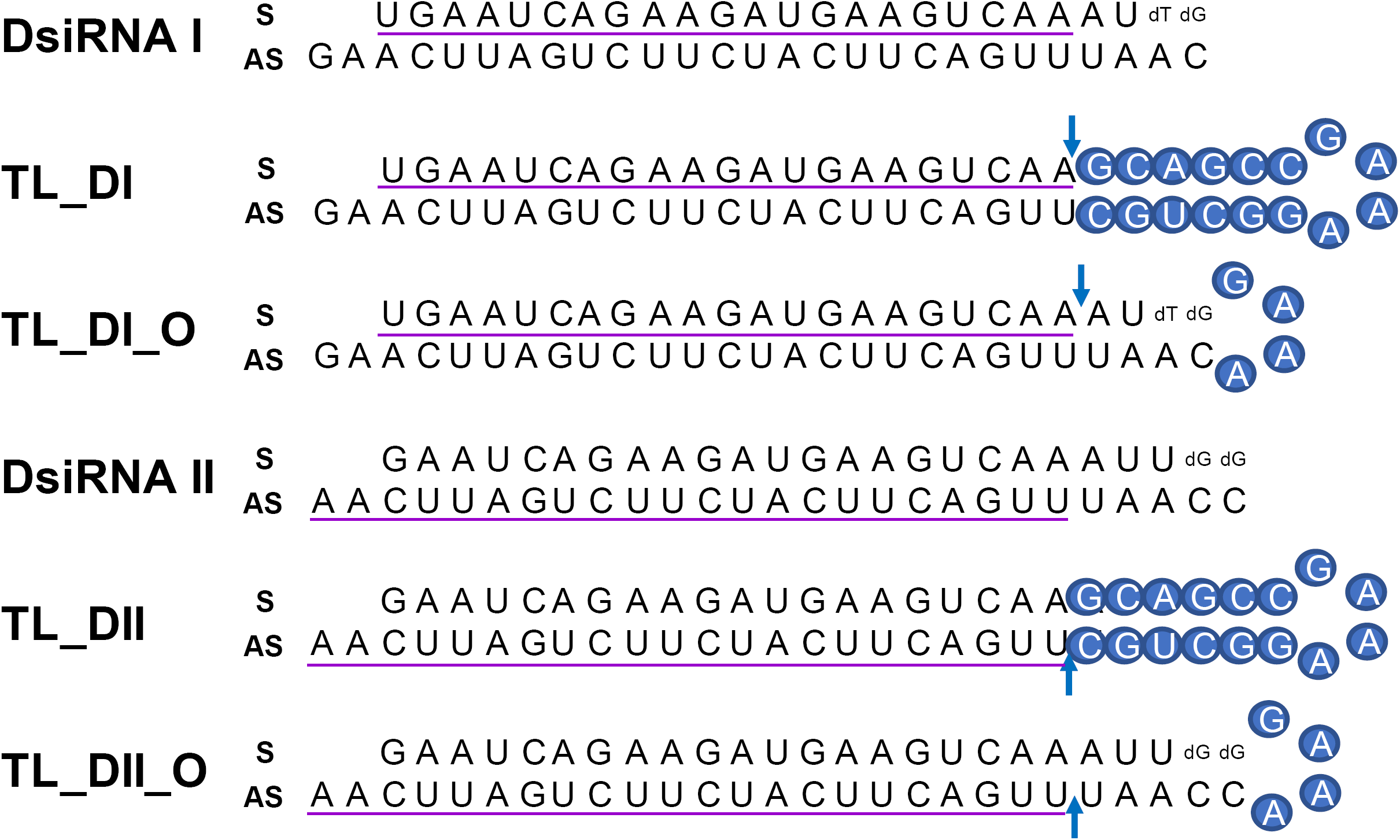
Structure of hnRNPH1-targeting DsiRNAs and tetra-looped DsiRNAs. We designed two DsiRNAs targeting hnRNPH1, DsiRNA I (**DI**) and DsiRNA II (**DII**), then designed tetra-looped versions. The tetra-looped DsiRNA I (**TL_DI**) has a 21-mer sense strand and 38-mer antisense strand, with a GC rich stem and 5’-GAAA-3’ tetra-loop (represented with blue circles). The tetra-looped DsiRNA I Original (**TL_DI_O**) has the same sequence as **DI**, with a 5’-GAAA-3’ loop sequence (blue circles). The tetra-looped DsiRNA II (**TL_DII**) has a 22-mer antisense strand and 36-mer sense strand, with a GC rich stem and GAAA tetra-loop (blue circles). The tetra-looped DsiRNA II Original (**TL_DII_O**) has the same sequence as DII, with a GAAA loop sequence (blue circles). The detailed sequences are listed in Table 1. Ribonucleotides are shown in upper case and deoxyribonucleotides as dN. The purple underlined sequence shows the active strand. The blue arrow indicates the nick between ribonucleotides.

### Generation of Dicer knockout HCT116 cells

We previously showed that chemically synthesized 25-to 27-nt-long DsiRNAs interacted with Dicer to facilitate the loading of small RNAs into RISC (27). DsiRNAs increased the gene silencing effect up to 100 times more efficiently than canonical siRNAs (17); recruiting the Dicer enzyme complex using DsiRNAs improves RISC assembly and gene silencing efficacy compared to 21-mer siRNAs. To investigate the effect of tetra-looped DsiRNAs on Dicer efficacy, we generated Dicer knockout cells by transfecting RNA-guided Cas9 endonuclease into the HCT116 cell line. We chose the HCT116 cell line because it is not aneuploid and often used for gene knockout studies. To knockout DICER in the human cell line HCT116, we used CRISPR/Cas9 technology. We designed guide RNAs complementary to the area near the genomic locus corresponding to the DExDc of DICER (Figure 2A and B; blue dot and letters). We included an indicator construct containing a green fluorescent protein (GFP) to show specific GFP expression in cells expressing Cas9. To generate a Dicer-deficient derivative of HCT116 cells, we isolated and expanded single-cell clones for further analysis. We obtained a series of three independent clonal cell lines (H2-1, H2-2, H2-3). After single-cell cloning, we purified genomic DNA (gDNA) and performed PCR using gDNA primers (Figure 2B, represented by green or orange arrows; Supplementary Figure 1). We mixed WT PCR product from parental HCT116 cells with Dicer H2 PCR product from single-cell clones of Dicer knockout HCT116 cells in equal quantities to produce heteroduplex molecules. We analyzed the cleavage products by DNA gel electrophoresis. By design, we expected the Surveyor nuclease enzyme to cleave the mismatched nucleotides in the heteroduplex molecule and generate two bands of 962 bp and 196 bp. However, the homodimer PCR products showed the undigested fragment. The Dicer H2-2 clone showed two bands in our electrophoresis assay (Figure 3). This suggested that Dicer H2-2 contained the mismatched nucleotides introduced by the CRISPR/Cas9 system, and it was selected for further analysis.

**Figure 2.**
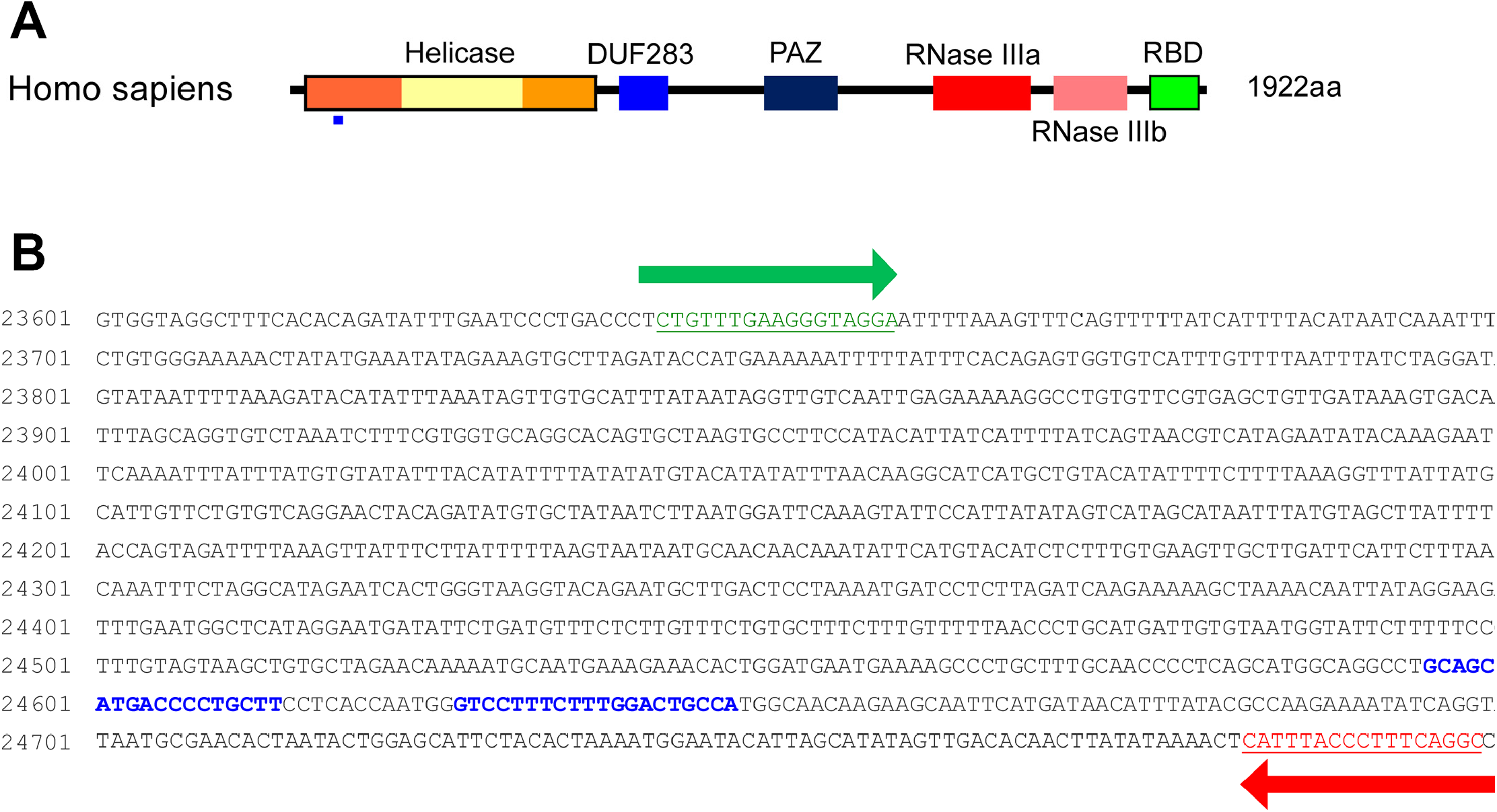
Dicer mutation by CRISPR/Cas9 system. A. Domain structure of Dicer protein. Blue rectangles indicate the regions corresponding to the genomic DNA (gDNA) sequences targeted by the CRISPR/Cas9 system. B: Sequence of Dicer gDNA. The two sites with blue letters show the sequence targeted by double-nickase CRISPR. Green and red arrows indicate primers for detecting Dicer gDNA.

**Figure 3.**
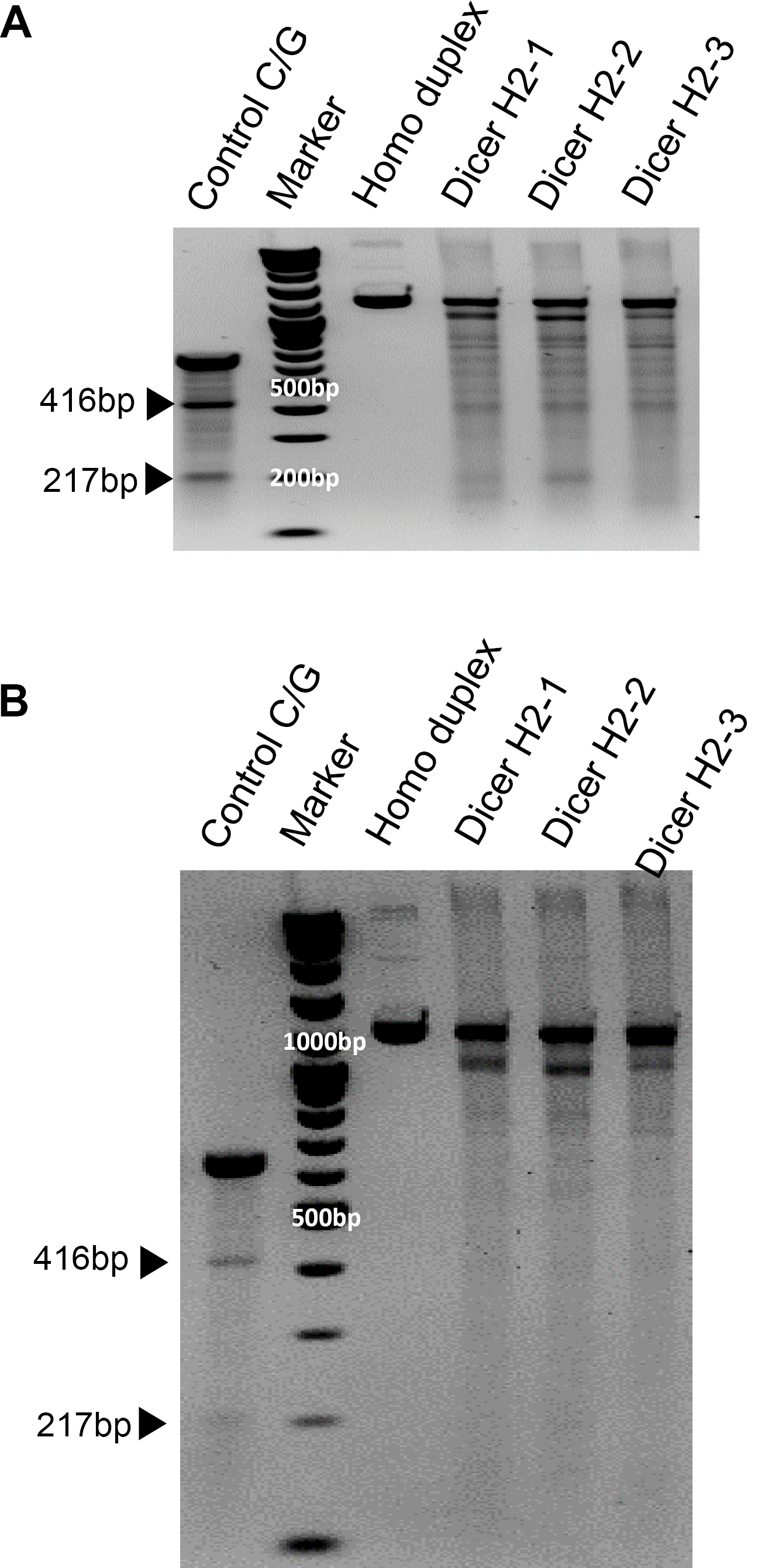
SURVEYOR assay comparing the efficiency of Cas9-mediated cleavage by double-nickase sgRNA in the human Dicer locus. DNA duplex formation and treatment with a nuclease. SURVEYOR assay gel showing a comparable modification of control G/C, which is 633 base pair (bp) Control DNA with a point mutation (Supplementary Figure 1B) bearing 416bp and 217bp. Homo duplex without mismatch did not cleave the nuclease, but hetero duplex (Dicer H2-1, H2-2, and H2-3) show the cleavage band. We run the gel short (A) and long (B) running. Arrowheads indicate cleavage products.

To confirm whether the mismatched nucleotides were included in the Dicer H2-2 clone, we analyzed the gDNA using Sanger sequencing. Using the CRISPR/Cas9 system, we designed two single-guide RNA (sgRNA; 20 bp: blue letters in Figure 4A) followed by a PAM sequence (orange letters in Figure 4A). We sub-cloned the PCR products, then sequenced each clone. All three clones had altered sequences (Figure 4B). Subsequent sequence analysis confirmed that the CRISPR/Cas9 system introduced DNA double-strand breaks at the target genomic sequences and thereby induced indels via error-prone nonhomologous end-joining repair (28).

**Figure 4.**
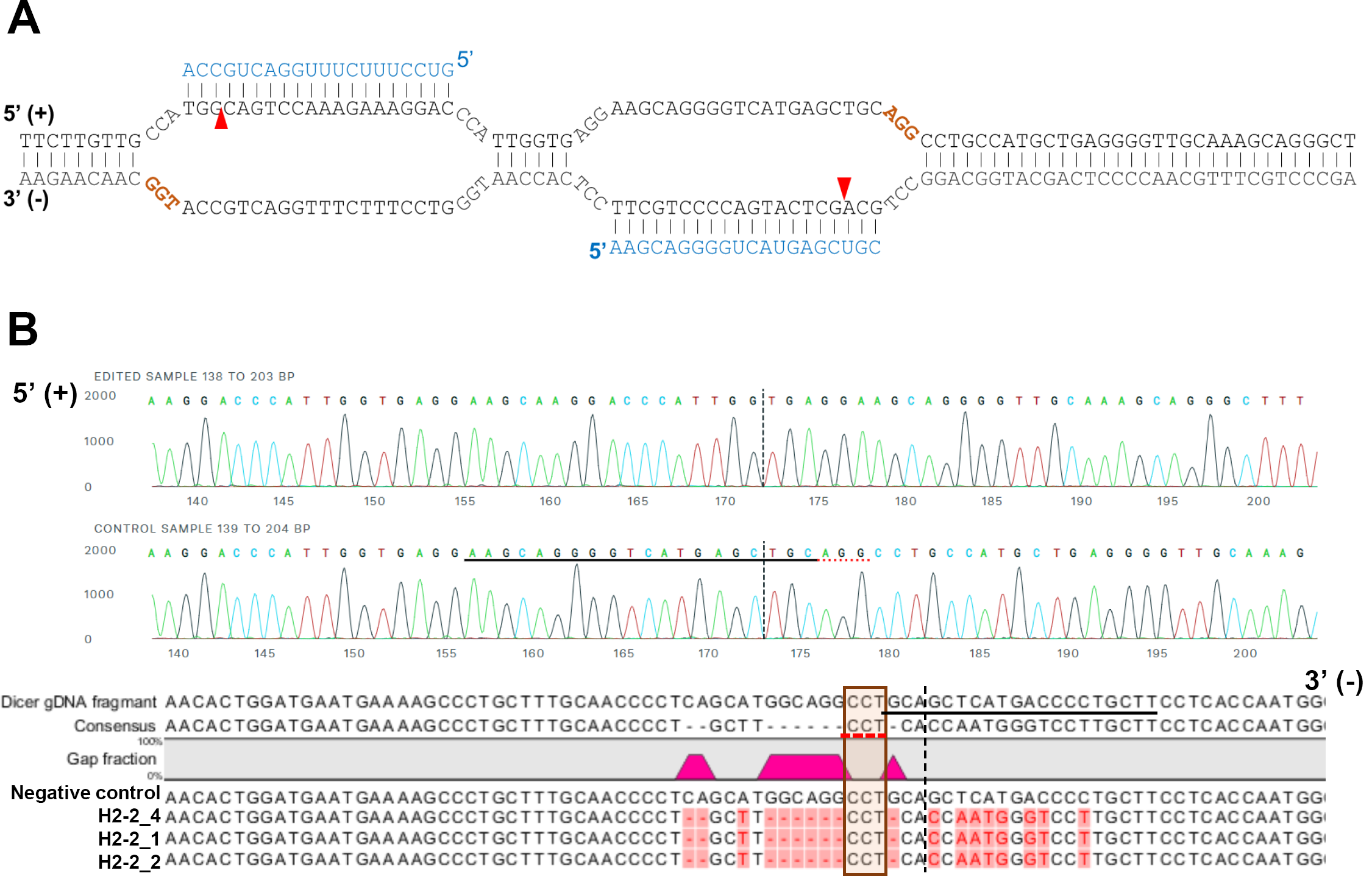
A representative sequence of the human DICER locus targeted by Cas9. A. Schematic illustrating DNA double nickase using a pair of sgRNAs guiding Cas9 nickase. Cas9 can cleave only the strand complementary to the sgRNA (Blue letters). A pair of sgRNA-Cas9 able to cleave Dicer gDNA. Expected cleavage site marks as Red arrow, respectively. sgRNA offset is characterized as the distance between the PAM (brown letters)- 5’-ends of the guide sequence of a given sgRNA pair. The brown letters are the PAM sequence. The red arrow is an expected cleave site by Cas9. B. Representative sequences of the genomic DNA in the helicase domain of human Dicer targeted by sgRNA. PAM area represents a red under-lined dot (upper) and boxed (bottom). The dotted vertical line is the expected cleavage site. We compared H2-2 clone sequence with the human Dicer genomic DNA sequence. The negative control is the gDNA sequence from WT HCT116.

### Tetra-looped DsiRNA enhanced siRNA efficacy

To investigate the efficacy of tetra-looped DsiRNA, we used dual-luciferase assays in WT HCT116 cells to separately detect the level of gene silencing conferred by the antisense or sense strand. We normalized the relative *firefly* luciferase activity of the DsiRNAs by dividing by *renilla* luciferase as an internal control, then converted 100 value of vector control. We first confirmed the suppression efficiency of DsiRNA in WT HCT116 cells. The DsiRNA I sense strand showed more significant suppression activity than the antisense strand (Figure 5A); in contrast, the DSiRNA II antisense strand was more effective than the DsiRNA II sense strand (Figure 5B). The inclusion of tetra-looped DsiRNAs did not change strand selectivity in WT HCT116 cells (Supplementary Figure 2). Next, we evaluated the silencing effect of each strand of tetra-looped DsiRNAs in WT HCT116 cells. The gene-silencing potency of tetra-looped DsiRNA significantly increased on the preferred strand for the DsiRNA I sense strand (Figure 5C) and DsiRNA II antisense strand (Figure 5F). However, RNAi activity was inconsistent on the non-preferred antisense strand of DsiRNA I and sense strand of DsiRNA II (Figure 5D and E). We designed a nick break between nucleotide bonds that we expected to be cut by Dicer and added tetra-loops to the DsiRNA. On the other hand, an intact guide strand is not necessary for cleavage by Dicer. The resulting increase in gene silencing activity of tetra-looped DsiRNAs might from RISC as a nick break on the guide strand makes it easily unwound. These findings suggest that the tetra-looped DsiRNAs are more efficient for gene silencing than the original DsiRNAs.

**Figure 5.**
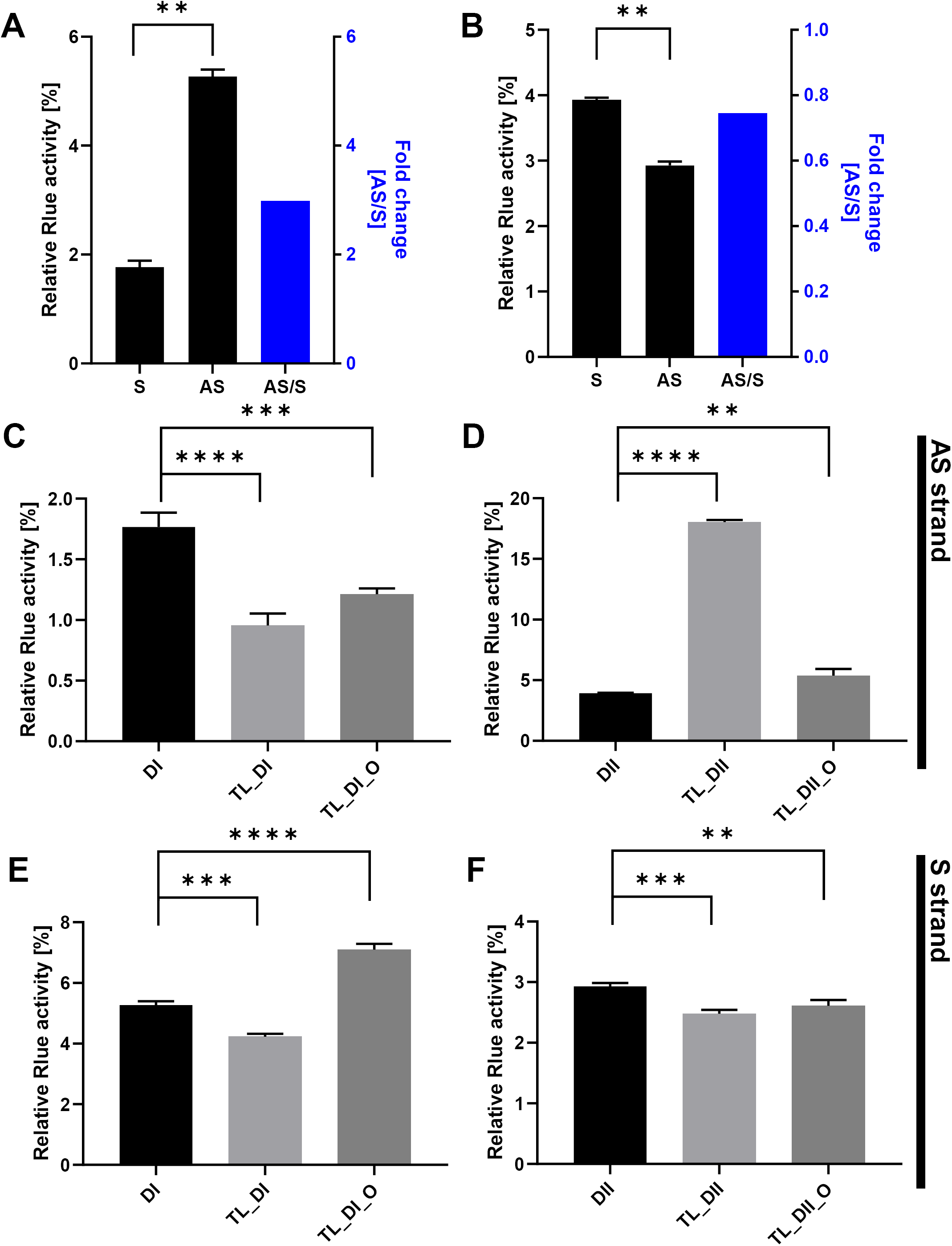
DsiRNAs efficacy in HCT116. Reporters contain the *Renilla* luciferase 3’ UTR of Antisense(AS_GS) or sense(S_PS) of hnRNPH1 transcript. Relative expression of the reporter with *Firefly* and *Renilla* luciferase determined by dual-luciferase assay. The internal control is the value of *firefly* luciferase. The *Renilla* activities were normalized to those of *firefly* and arbitrarily set at 100. Mean values and standard deviations from three independent experiments are shown. (student t-test *P<0.05, **P<0.01, ***P<0.001, and ****P<0.0001) We transfected dual-luciferase reporter and variant DsiRNAs into HCT116. The DsiRNA represent A. DsiRNA I, B. DsiRNA II, C: DsiRNA I_S, D: DsiRNA II_S, E: DsiRNA I_AS, and F:DsiRNA II_AS.

### Gene silencing activity is controlled by Dicer

To confirm that Dicer is the main protein involved in small RNA biogenesis, we transfected the DsiRNAs and tetra-looped DsiRNAs into WT HCT116 cells and H2-2 Dicer knockout HCT116 cells. We conducted luciferase assays to quantitate gene silencing. If DsiRNA-induced gene silencing does not occur through Dicer, we would expect equivalent levels of luciferase activity in both WT HCT116 and H2-2 Dicer knockout cells. However, all strands of DsiRNAs, including tetra-looped DsiRNAs, showed significantly reduced gene silencing activity in H2-2 Dicer knockout cells compared to WT HCT116 cells (Figure 6). These results show that Dicer processing is essential to small RNA-mediated gene silencing of both DsiRNAs and tetra-looped DsiRNAs.

**Figure 6.**
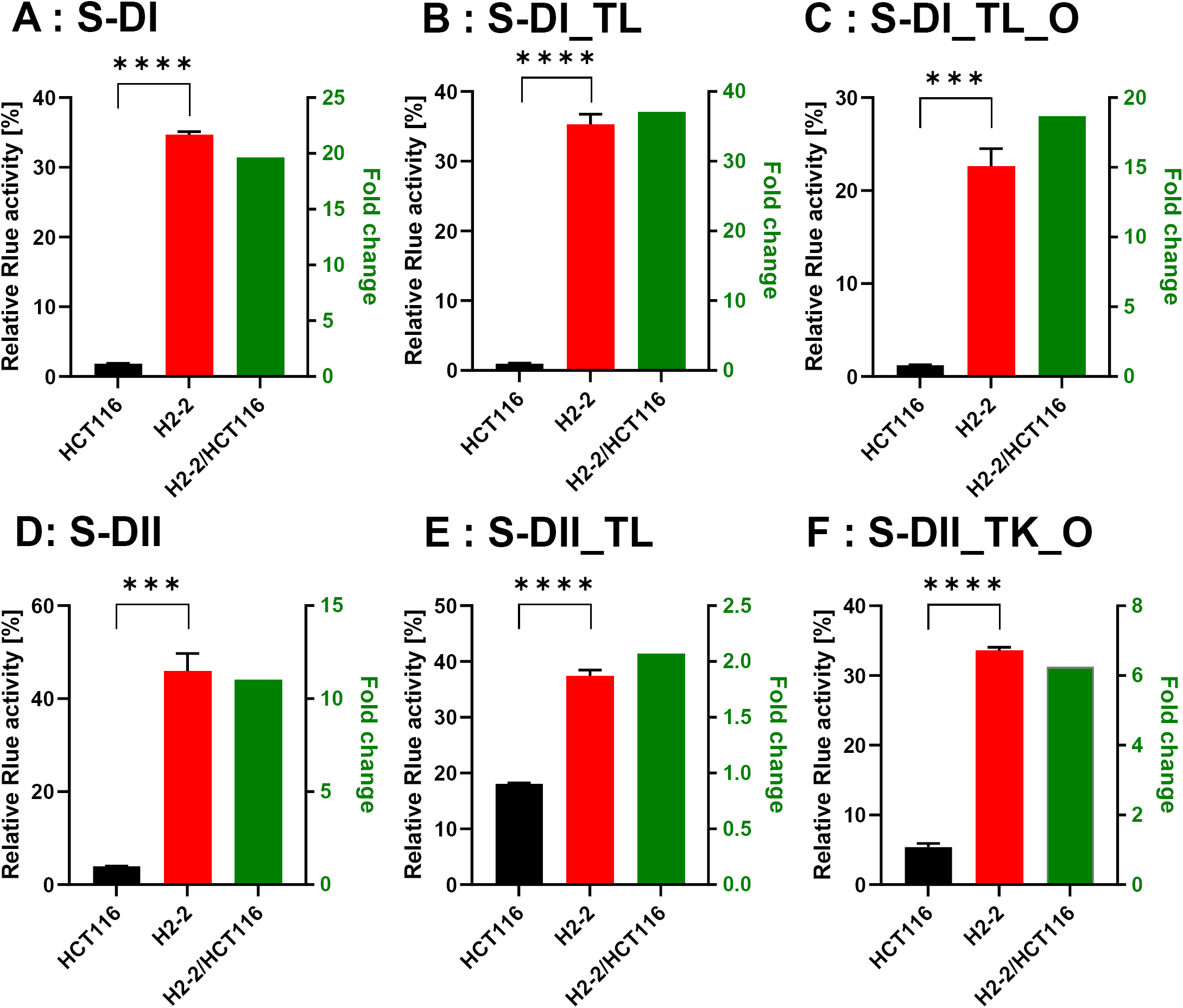
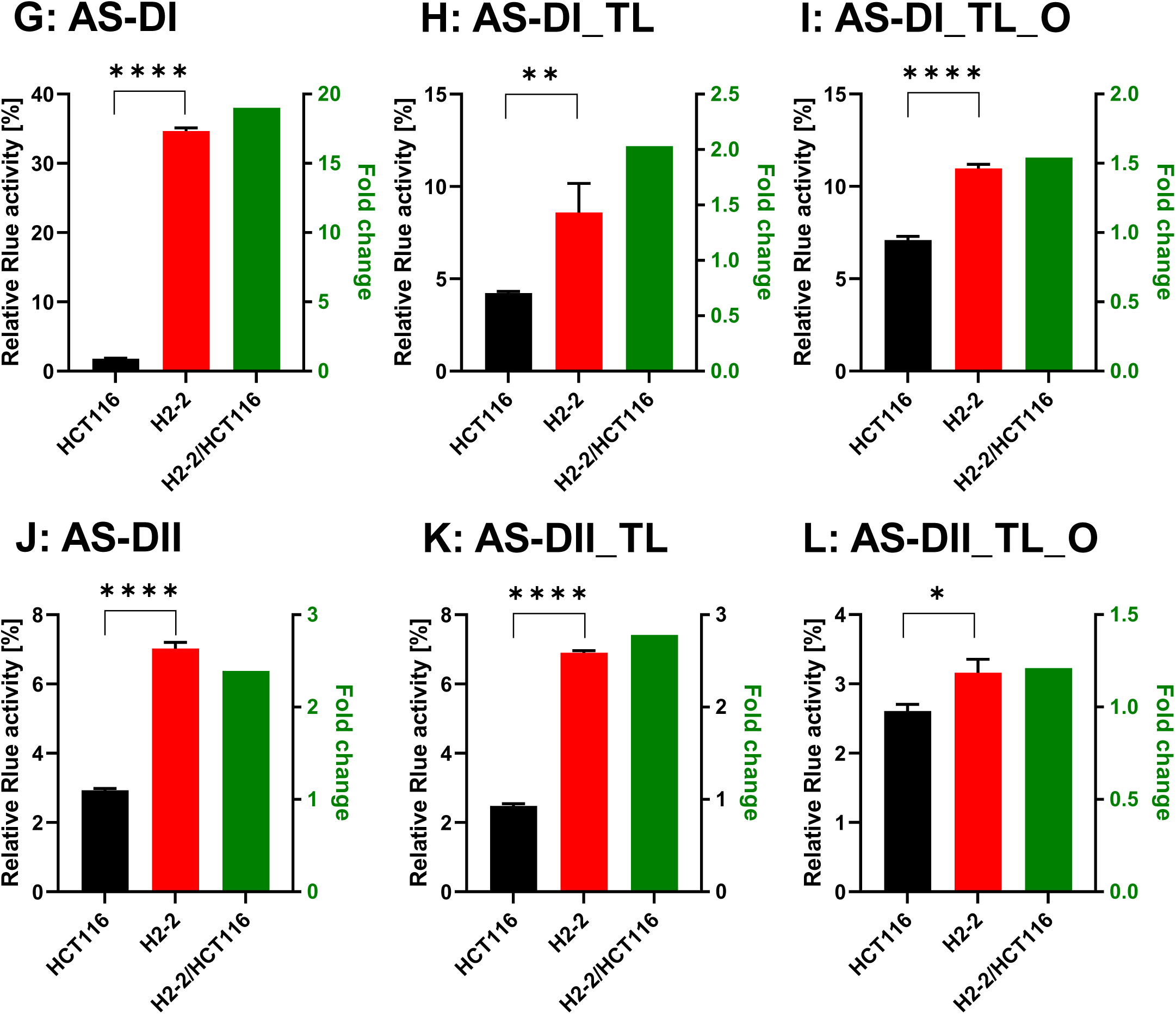
Compare with DsiRNAs efficacy in HCT116 and H2-2. The detection method was the same as Figure 6. We transfected the dual-luciferase reporter (S: vector with targeting sense strand of DsiRNAs, AS reporter with targeting antisense strand of DsiRNAs) and DsiRNAs into HCT116 and H2-2 (Dicer knockout HCT116: red bar). We detected the fold decrease of gene silencing efficiency in H2-2A to calculated the activity of H2-2 divided by HCT116 (green bar). It represented the right *y*-axis. The graphs showed the gene silencing activity of A: sense strand of DI (S-DI), B: sense strand of DI_TL (S-DI_TL), C: sense strand of DI_TL_O (S-DI_TL_O), D: sense strand of DII (S-DII), E: S of DII_TL (S-DII_TL), F: S of DII_TL_O (S-DII_TL_O), G: antisense strand of DI (AS-DI), H: antisense strand of DI_TL)AS-DI_TL), I: antisense of DI_TL_O (AS-DI_TL_O), J: antisense of DII (AS-DII), K: antisense of DII_TL (AS-DII_TL), and L: antisense of DII_TL_O (AS-DII_TL_O). Data represent the mean ± S.D. of three independent repeats. An asterisk indicates statically significant differences at the level of *P<0.05, **P<0.01, ***P<0.001, and ****P<0.0001 tested by student’s t-test.

## Discussion

In this study, we evaluated the gene silencing activity of DsiRNAs and tetra-looped DsiRNAs in WT and H2-2 HCT116 cells to better understand Dicer mechanisms. We found that the preferred strand of tetra-looped DsiRNA improves RNAi gene silencing activity (Figure 5). Previously, shRNA, which has a stem-loop structure, was shown to induce stable and long-lasting gene silencing activity (29-31). The RNAi gene silencing activity of shRNAs with stem-loop structures requires Dicer activity (23), possibly because the terminal loops of shRNA and pre-miRNAs share similar RNA structures, which interact with the N-terminal helicase domain of Dicer (25,32-34). Our results suggest that tetra-loop structures are involved in essential Dicer ribonuclease functions, and may have potential alternative effects through interaction with RNA and protein. Other studies of Dicer and pre-miRNA crystal structural analysis revealed that the N-terminal domain of Dicer binds the more stable region of single-stranded hairpin looped pre-miRNA (35-37). The N-terminal domain of Dicer is bound to TRBP, which advances the RNA-binding affinity of Dicer (38) and reinforces cleavage accuracy (39,40). We predict that Dicer possesses not only ribonuclease activity but can also affect RNA stability and that TRBP is an RNA cofactor for Dicer complexes, such as for DsiRNA with tetra-loops, which can increase gene silencing activity. This suggestion agrees with previous reports indicating that Dicer can globally bind to stem-looped RNAs without cleavage activity and influences the fate of targeted transcripts by gene silencing (41). Based on the experimental data presented here, DsiRNAs with added tetra-loops showed higher efficiency in gene silencing, but this activity disappeared in Dicer knockout HCT116 cells (Figures 5 and 6). This finding suggests a potential complex in which the stem-loop structure of small RNA and Dicer can improve gene silencing activity.

Dicer is an essential protein for small RNA biogenesis and has recently been reported to act as a multifunctional protein which is not limited in miRNA and siRNA (42). The processing of long transfer RNAs (tRNAs) to small RNAs, broadly termed tRNA fragments (tRFs), is dependent on Dicer (43-45). Sno-derived RNAs (sdRNAs), which are derived from small nucleolar RNAs (snoRNAs), showed reduced expression in Dicer mutants (46,47). The depletion of most miRNA species is detected following Dicer ablation (48). Our study provides evidence that tetra-loop DsiRNAs exhibit more potent gene silencing that depends on Dicer (Figure 6). We thus speculate that Dicer is not only identified by its endonuclease activity for small RNAs but also can be stably bound with tetra-looped DsiRNA to enhance gene silencing activity.

Our observations raise intriguing questions regarding the mechanism of how Dicer improves the activity of RNA gene silencing. Interestingly, Dicer can bind to many classes of RNA molecules, in which the interaction does not necessarily lead to dicing. Specific stem-loop structures that bind with Dicer have been identified by human transcriptome-wide analysis and are called “passive Dicer-binding sites” (41). Dicer is believed to participate in the correct guide strand for Ago2 loading to generate RISC (49,50). Our finding suggests that Dicer can generate its own siRNA and can function to stabilize small RNA. We conclude that Dicer-mediated processing of tetra-looped DsiRNAs subsequently facilitates a more stable interaction and improved efficiency.

## Experimental precedures

### DsiRNA duplex

All RNA strands for DsiRNAs were synthesized as single-strand RNAs (ssRNAs) by Integrated DNA Technologies with high-performance liquid chromatography purification and resuspended in RNase-free water (Table 1). ssRNAs were annealed to form DsiRNA duplexes at 95°C for 5 minutes, then incubated for 4 hours at room temperature, before being aliquoted (10 µL in 1.5-mL tube) and stored at - 80°C.

### Cell culture and transfection

HCT116 cells were cultured in DMEM supplemented with 10% fetal bovine serum and penicillin-streptomycin at 37°C in 5% CO_2_ with humidification. HCT116 cells were transfected with Lipofectamine 2000 (Life Technologies). The Dicer knockout HCT116 cells were generated by employing CRISPR technology (DICER Double Nickase plasmid h2, Santa Cruz, sc-400365-NIC-2 following the manufacturer’s protocol. The DICER genome sequence and sgRNA targets are represented in Figure 2.

### Dual-luciferase assay

To generate the reporter plasmids psi-hnRNPH-S (sense reporter) and psi-hnRNPH-AS (antisense reporter), a 343-bp PCR fragment of hnRNPH cDNA (Acc.: NM_005520) was cloned in the 3′-UTR of the humanized *Renilla* luciferase gene in the psiCHECKTM-2 vector (Promega) in either the sense or antisense orientation. The cells were co-transfected with 0.1 µg of the plasmid with the dual-luciferase reporter system and DsiRNAs using 2 µl of Lipofectamine 2000 in a 48-well plate. Luciferase assays were performed 48 hours after transfection, using the dual-luciferase reporter assay system (Promega). Firefly Luciferase activity was normalized to the Renilla Luciferase activity and then to its own control, the activity of which was set to 100.

### Surveyor nuclease assay

We purified gDNA from CRISPR/Cas9-treated HCT116 cells to confirm Dicer knockout. We amplified segments containing the sgRNA target site by PCR, using 100 ng gDNA with a representative primer shown in Figure 2B. We purified the DNA from PCR products, then mixed 300 ng PCR products obtained from WT and Dicer knockout mutant HCT116 cells, and denatured them by heating at 99°C for 5 min in a thermocycler. We then formed heteroduplexes and homoduplexes by cooling down to room temperature.

We performed the Surveyor nuclease assay using a Surveyor^®^ mutation detection kit (IDT). We mixed each sample with 1 µL of Surveyor Enhancer S, 1 µL of SURVEYOR nuclease S, and 4 µL of 0.15 M MgCl2 in a 50-µL reaction. We incubated the mix for 60 min at 42°C and stopped the reaction by adding 4 µL of the stop solution provided in the kit. The reactions were either kept at −20° or used immediately for electrophoresis.

### Statistical analysis

Statistical analyses were performed using Student’s t-test. All data represent the mean ± SD of at least three independent experiments.

## Footnotes

This study was supported by Cancer Center Support Grant 033572 and National Institute of Health AI042552 (to J. J. R.). The authors declare that they have no conflicts of interest with the contents of this article.

## The abbreviations used are

DsiRNA: Dicer-substrate siRNA
rRNA: ribosomal RNA
dsRNAs: double-stranded RNAs
siRNAs: small interfering 21-22bp RNAs
RISC: RNA-induced silencing complex
miRNA: microRNA
pri-miRNA: hairpin-containing primary transcripts
shRNAs: short hairpin RNAs
DExD: DEAD-like helicases domain
TRBP: transactivation response element RNA-binding protein
WT: wild-type
RNAi: RNA interference
hnRNPH1: heterogeneous nuclear ribonucleoprotein H
TL: tetra-loop
GFP: green fluorescent protein
gDNA: genomic DNA
sgRNA: single-guide RNA

## Figure Legend

**Supplementary Figure 1.**
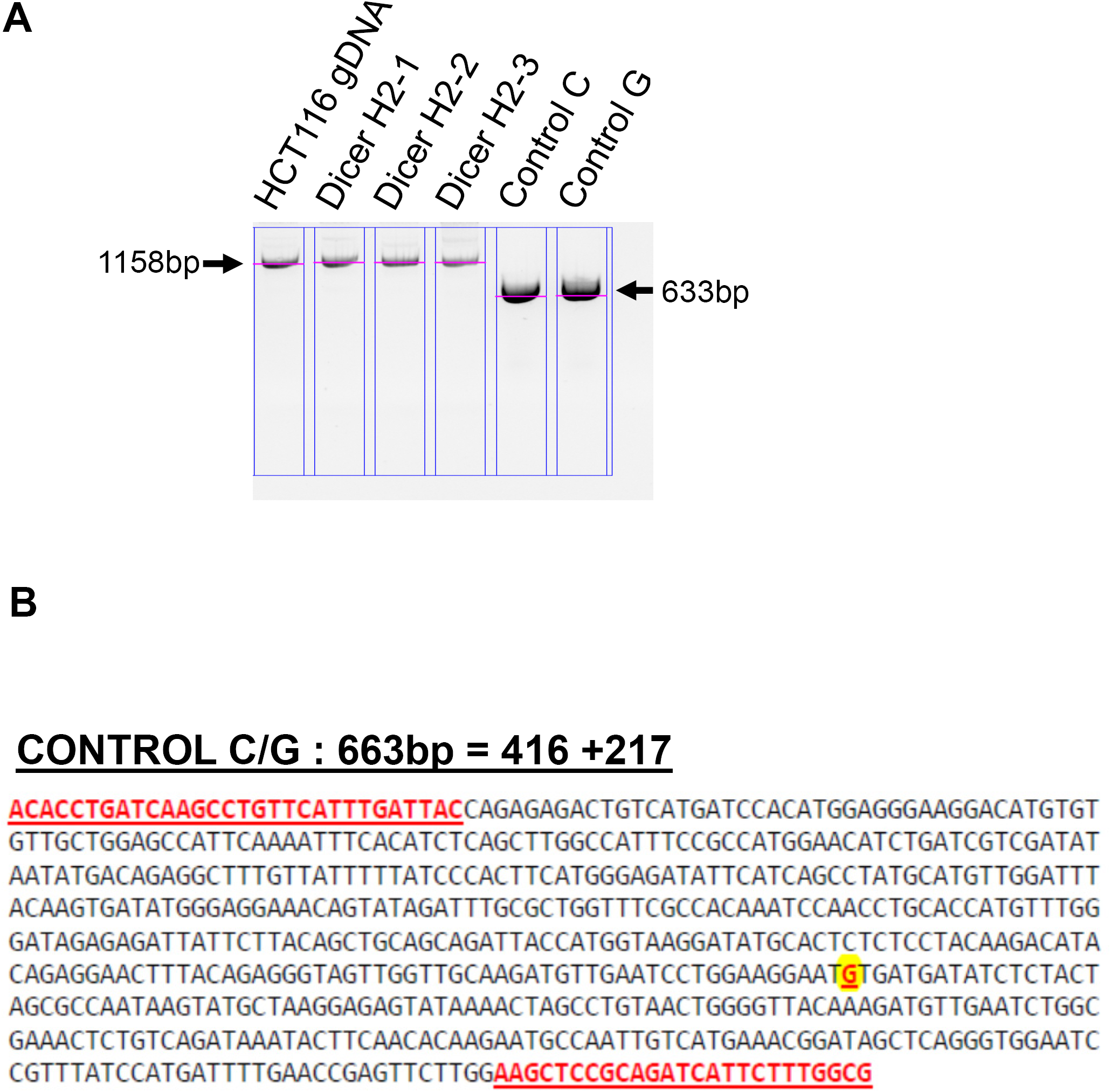
Genomic DNA PCR from single-clone cells. A.PCR using genomic DNA for single cell clone of H2-1, H2-2, and H2-3 (1158bp), control C, and control G (633bp). B. The sequence presents the control DNA sequence. Yellow marked G is point mutation site, it is G or C. Red letter and under line is forward and Reverse primer site for gDNA PCR

**Supplementary Figure 2.**
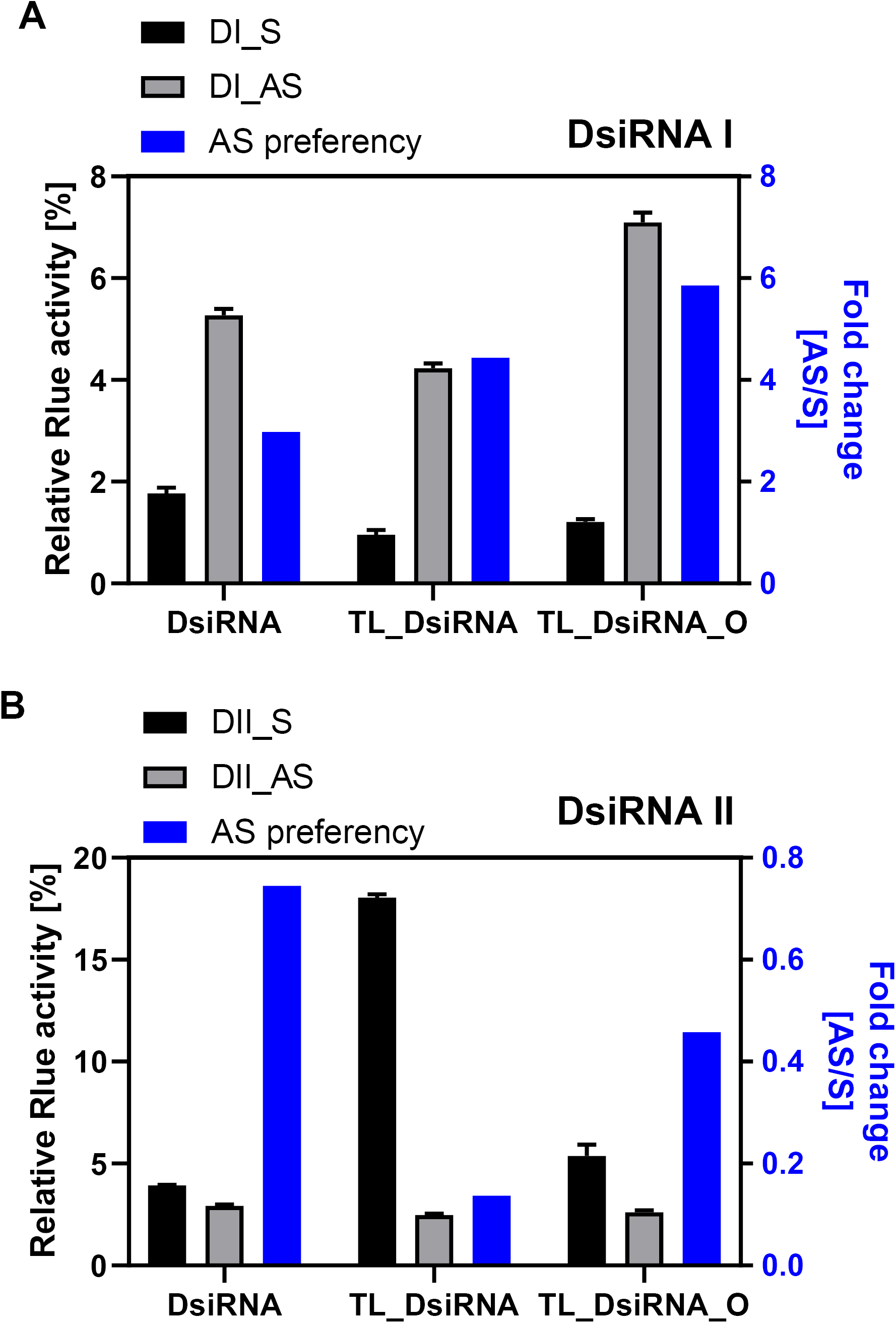
Gene silencing activity in each strand of DsiRNAs. The version of DsiRNA I(A) and DsiRNAII(B) transfected HCT116 cells were lysed in 1X passive lysis buffer, and dual-luciferase activities were determined. We calculated the AS preference for Relative Rlue activity of AS divided by S activity (AS preferency: blue bar). DsiRNA I showed more 2-fold AS preferency. In contrast, the AS preferency of DsiRNA II below 1. DsiRNAIIs have preferred the S strand in DsiRNA II, TL-DsiRNAII, and TL-DsiRNAII-O. The data shown represent the means of three experiments, with the bar showing SD.

